# Intra-annual precipitation variability mutes or magnifies the impact of wet and dry years on plant biomass

**DOI:** 10.64898/2026.02.03.703587

**Authors:** Tyson J Terry, Steven Higgins, Alexandra Hamer, Peter B. Adler, Jonathan D. Bakker, Lars A. Brudvig, Elizabeth T. Borer, Miguel N. Bugalho, Maria C. Caldeira, Jane A. Catford, Qingqing Chen, Scott L. Collins, Chris R. Dickman, Nicole Hagenah, Kimberly Komatsu, Johannes M.H. Knops, Yujie Niu, Xavier Raynaud, Anita C. Risch, Eric W. Seabloom, Glenda Wardle, Jenifer L. Yost, Anke Jentsch

**Affiliations:** School of Life Sciences, Arizona State University, Tempe, USA, 85282; Chair of Disturbance Ecology, Bayreuth University, Bayreuth, DE 95447; Chair of Plant Ecology, Bayreuth University, Bayreuth, DE 95447; Centre for Alternative Technology, Llwyngwern Quarry, Pantperthog, Machynlleth, UK SY20 9AZ; Department of Wildland Resources and the Ecology Center, Utah State University, Logan, UT 84322, USA; School of Environmental and Forest Sciences, University of Washington, Seattle, WA 98195 USA; Department of Plant Biology and Program in Ecology, Evolution, and Behavior, Michigan State University, East Lansing, MI 48824 USA; Department of Ecology, Evolution, and Behavior, University of Minnesota, St. Paul, MN 55108 USA; Centre for Applied Ecology “Prof. Baeta Neves” (CEABN-InBIO), School of Agriculture, University of Lisbon, Portugal; Forest Research Centre (CEF), TERRA Associated Laboratory, School of Agriculture, University of Lisbon, Portugal; Department of Geography, King ‘s College London, 40 Aldwych, London, WC2B 4BG, UK; Fenner School of Environment & Society, The Australian National University, Canberra, ACT 2600, Australia; Institute of Ecology and Evolution, University of Bern, Baltzerstrasse 6, 3012 Bern, Switzerland; Senckenberg Museum of Natural Sciences, Görlitz, Germany; Department of Biology, University of New Mexico, Albuquerque, NM 87131 USA; Desert Ecology Research Group, School of Life and Environmental Sciences, The University of Sydney, NSW 2006, Australia; Mammal Research Institute, Department of Zoology and Entomology, University of Pretoria, Pretoria, South Africa; School of Life Sciences, University of KwaZulu-Natal, Private Bag X01 Scottsville, Pietermaritzburg, 3209, South Africa; Department of Biology, University of North Carolina at Greensboro, Greensboro, North Carolina 27402, USA; Health & Environmental Sciences Department, Xi’an Jiaotong-Liverpool University, Suzhou, Jiangsu, China; Institut d’écologie et des sciences de l ’environnement de Paris, Sorbonne Université, Université Paris Cité, Univ Paris Est Créteil, CNRS, IRD, INRAE, IEES, F-75005 Paris, France; Swiss Federal Institute for Forest, Snow and Landscape Research WSL, Zuercherstrasse 111, 8903 Birmensdorf, Switzerland; Desert Ecology Research Group and School of Life and Environmental Sciences, ARC Training Centre in Data Analytics for Resources and Environments (DARE), The University of Sydney, Sydney, NSW, Australia; USDA-ARS Grassland Soil and Water Research LaboratoryTemple, TX, 76502, USA

## Abstract

As the atmosphere warms, both the amount and timing of precipitation are changing, but how those two trends interact to influence plant growth remains unresolved. Using 5-15 years of data from 48 globally distributed grassland sites, we quantified how the temporal distribution of precipitation within the year (evenness) interacts with annual precipitation amount and nutrient limitation to influence plant biomass. Annual precipitation anomalies had a large influence on plant biomass when intra-annual precipitation was more evenly distributed (frequent small events) and little impact when less even (infrequent large events). This relationship was consistent across aridity gradients and nutrient limitations and strongest in systems with warm wet seasons. Our work shows that the response of plant systems to changes in annual precipitation amount are largely dependent on how evenly it is temporally distributed.

## Main Text

Global shifts in precipitation consist of changes in annual means, interannual variability, and the temporal distribution of precipitation ^1–3^. Variation in annual precipitation amount can exacerbate or relieve water limitations for agriculture ^4^, human water supply ^5^, and plant growth ^6^. Intra-annual precipitation distribution, in particular how evenly total precipitation is distributed within the year (precipitation evenness), also has potent effects on ecosystems ^7,8^ and society ^9^ with anomalies tied to flooding and prolonged dry periods between rainfall events ^10,11^. The temporal packaging of precipitation combines with total annual precipitation to determine the duration and quantity of water in soils ^7,12,13^. Precipitation trends and projections indicate increasing variability in annual amounts and less even precipitation as the warming atmosphere both demands and releases larger quantities of water ^1–3^. Larger and less frequent rain events can lead to deep water infiltration or high erosion depending on soil texture and water saturation ^12,14^, and may also create periods of dry soil and water-stress for plants between precipitation events if precipitation amounts do not match evaporative demands ^11,15^. Furthermore, intra-annual precipitation distribution may affect nutrient availability through effects of soil drying-rewetting on soil microbial activity ^16–18^, potentially extending the impact of precipitation variability beyond its direct effects on plant-available water.

Efforts to synthesize the independent effects of annual precipitation amount and precipitation evenness on plant growth have revealed large spatial and temporal inconsistencies in effect size ^1,6,19^, indicating the omission of important factors that modify the effect of precipitation amounts on plant growth. Currently, most approaches do not model the interaction between annual precipitation amount and evenness due to the use of evenness metrics, such as mean precipitation frequency and/or event size, that are strongly correlated with annual precipitation amount ^20,21^. Use of these metrics not only make it statistically difficult to separate effects of precipitation amount from those of evenness on plant growth ^1^, but also limit inference by isolating precipitation inputs (event size) from their temporal distribution (event frequency).

Precipitation evenness changes nutrient availability with accumulations or declines occurring during dry periods or deluge ^7,22^, which may then alter subsequent plant growth when water becomes available. However, the influence of evenness on nutrient limitations for plant growth remains poorly understood, as studies generally manipulate precipitation evenness and nutrients independently, without investigating their interactions. The complicated interplay of these factors stresses the need for a controlled cross-site analysis with methodology that allows investigation of precipitation evenness that occurs alongside variability in precipitation amount and nutrient availability.

Here we investigated the independent and interactive effects of precipitation amount and evenness on plant biomass. We utilized grassland sites as they represent an abundant ecosystem that is known to be responsive to fluctuations in annual precipitation amounts^6,23^. Our sites occur across an aridity gradient along with adjacent nutrient addition plots to determine if interactive effects of evenness and amount decline with water availability and if they are driven by impacts of evenness on nutrient limitation. We used an unranked Gini index (UGI) to quantify precipitation evenness ^3,24^(Fig. 1). The UGI measures how unevenly rainfall is distributed across time by measuring the deviation of the observed cumulative rainfall accumulation over time compared to a cumulative distribution that would arise if rainfall event sizes and frequency are evenly distributed over the same period (Fig. 1). An unranked Gini index allows the size and timing of a precipitation event to provide different impacts whether it happened after a dry period or after a period of substantial precipitation. Precipitation evenness is uncorrelated with precipitation amount (R_spearman_ = 0.03, Fig S5) because it is calculated using cumulative percentages of total annual precipitation rather than metric amounts, allowing us to investigate the independent and interactive effects of precipitation amount and evenness on plant growth.

**Figure 1.**
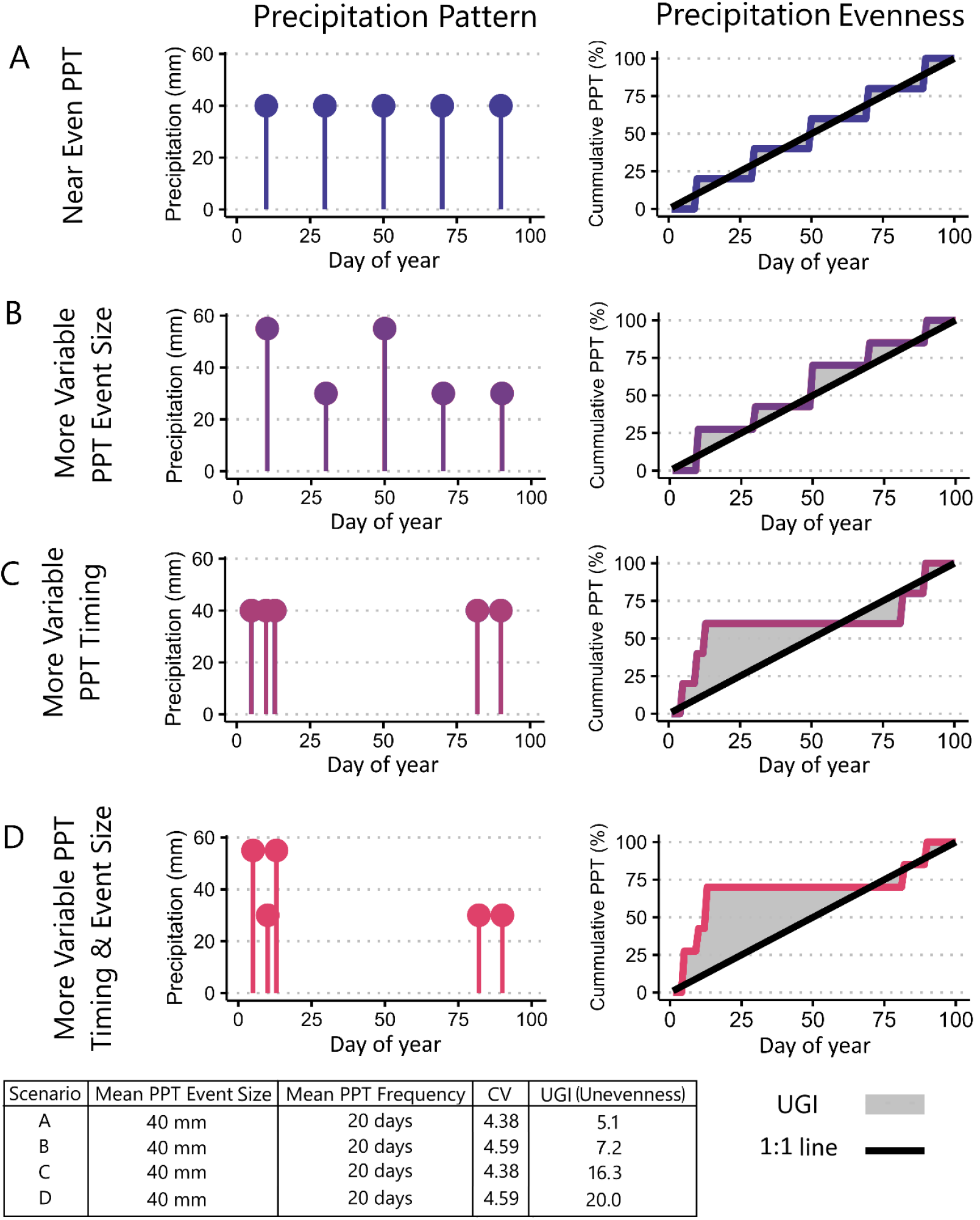
Hypothetical patterns of intra-annual precipitation events (panels A-D) and their associated variability metrics. Total precipitation (PPT) amounts are held constant across 100 days in all scenarios. Mean precipitation event size and frequency are calculated as the mean of all non-zero precipitation events and the mean days between non-zero precipitation events. Precipitation evenness is shown in the line graphs on the right and is quantified by the unranked Gini index (UGI) which corresponds to the annual absolute difference between the black 1:1 line and the colored line (total grey area).

We analyzed 5-10 years of annual plant biomass from 48 grassland sites across 6 continents within the Nutrient Network (www.nutnet.org) to test the following hypotheses: 1) Plant biomass responses to annual precipitation amounts are regulated by evenness as dry period length and infiltration of water into soils modify the temporal availability of annual precipitation amounts for plant growth. 2) The strength of the interaction will decline at sites with high mean annual precipitation and in nutrient addition plots due to the diminished impact of precipitation evenness on water and nutrient limitations for plant growth.

We found that evenness clearly mediated the impact of precipitation amount on plant biomass (Fig. 2). The impact of precipitation amount on plant biomass was highest when precipitation evenness was high and was insignificant under low evenness (Figs. 2 and S2). Likewise, with a mixed effects approach that accounts for additional spatial variation among sites, such as MAP and mean annual air temperature (MAT), we still found a significant interactive effect of precipitation evenness and precipitation amount (Fig, 3, t = -2.090, P = 0.037) on plant biomass. However, we didn ‘t observe any direct effects of evenness on plant biomass (t = 0.237, p = 0.813). This validates previous studies that found high sensitivity of plant growth to shifts in temporal precipitation variability ^19^, and explains why the effects of precipitation evenness are not independent from the effects of precipitation amount (Figs. 2 and 3).

**Figure 2.**
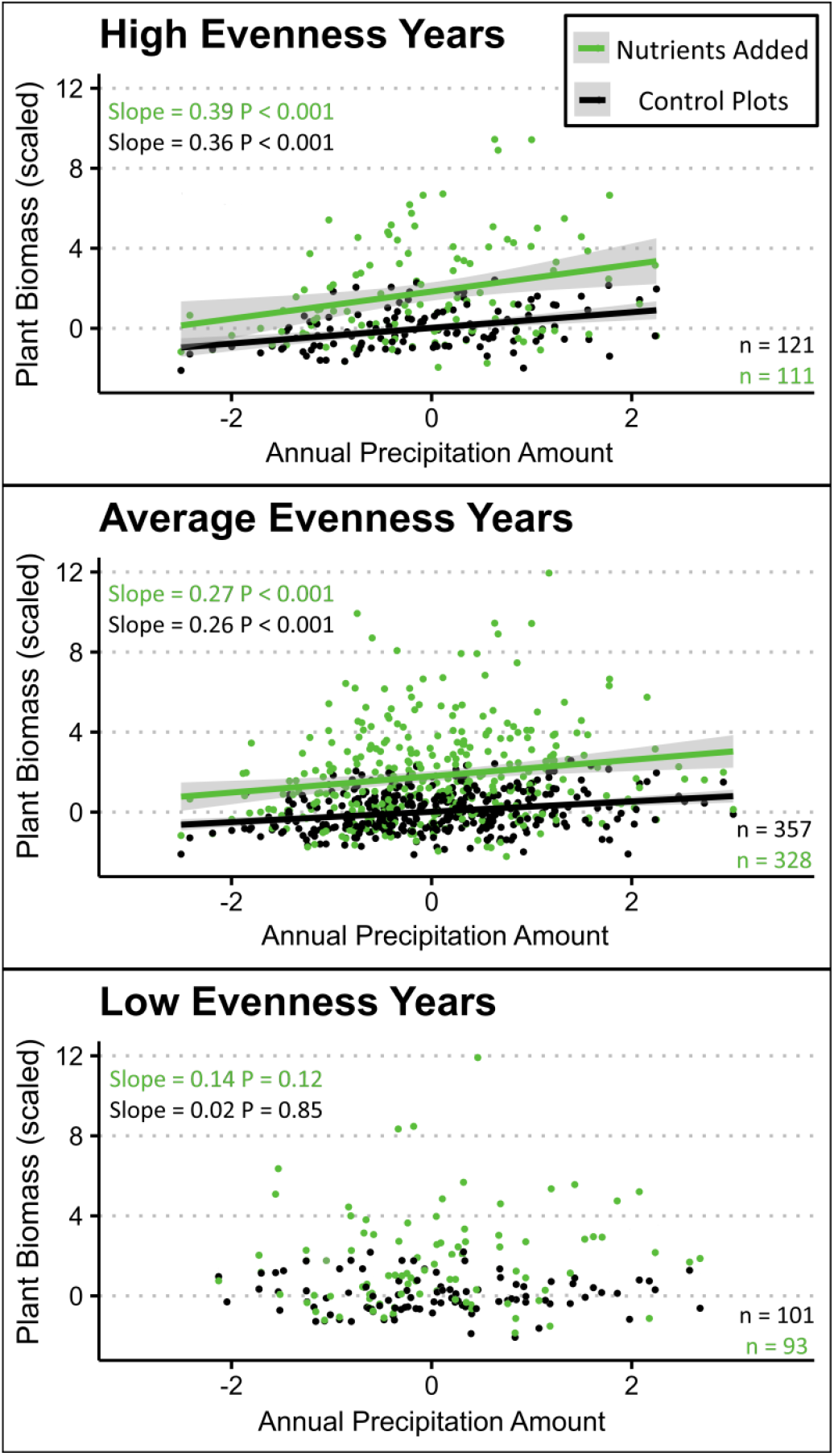
The effect of annual precipitation amount on plant biomass is most apparent under highly even intra-annual precipitation regardless of nutrient addition. Points represent shifts in total live biomass relative to site means in the control plots across all years. Lines represent linear regressions with their respective slope, significance, and number of data points listed in text labels. Solid lines indicate significant (P<0.05, two tailed p value) relationships, while dashed lines represent insignificant relationships. Annual precipitation (precipitation) amounts are annual precipitation totals scaled by site-specific means and standard deviations (2004-2023). High and low evenness categories represent years with evenness values greater or less than one standard deviation, respectively, from site-specific mean evenness. Categories of precipitation evenness were selected for display purposes only; all statistical analysis using a continuous variable.

**Figure 3.**
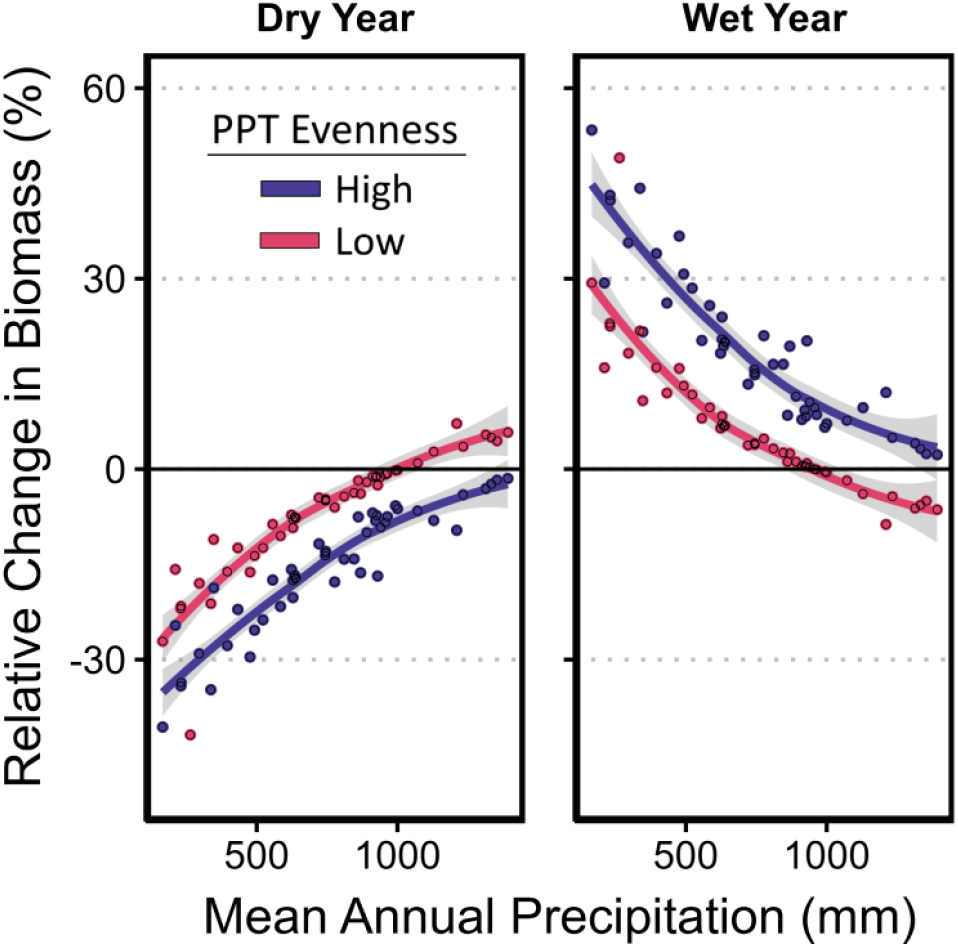
The effect of wet and dry years on plant biomass varies across a mean annual precipitation gradient but the effect of evenness remains consistent. Lines and points represent relationships and site level predictions from our model (R_sqr_ = 0.73). The light grey error margin represents the 95% confidence interval for model predictions. Dry and wet year scenarios are defined as years with one standard deviation above or below the site-specific average annual precipitation amount (2004-2023). High and low evenness scenarios represent values of one standard deviation above or below the site-specific evenness average (2003-2023). Categories of precipitation evenness and wet/dry years were selected for display purposes only; all statistical analysis using a continuous variable.

Less even precipitation increased expected plant biomass during dry years and decreased expected plant biomass during wet years (Figs. 2 and 3). This pattern has been observed in arid environments where soils are generally coarse and high evaporative demand reduces the benefits of small and frequent precipitation events (high evenness) for plant growth ^7,25^. However, we were surprised to find support for this pattern in a dataset including sites with high MAP (600-1200 mm). We hypothesize that large precipitation pulses during wet years led to high runoff and/or introduced periods of dryness that negated the benefit of above-normal precipitation years. The potential for runoff and erosion is highest when soil is saturated, or water inputs exceed infiltration rates ^26^. Less even precipitation during years with high precipitation amounts may saturate the rooting zone and lead to high runoff and the loss of water ^25,26^ and nutrients from the local system. Less even precipitation also creates post-precipitation dry periods with little cloud cover ^27^, increases soil water deficit ^28^, and raises atmospheric vapor pressure deficit ^29^ that may reduce the benefit of additional precipitation during wet years.

A three-way interaction of annual precipitation amount, precipitation evenness, and MAP on plant biomass was not significant in our linear regression model (t = -0.201, p= 0.84), indicating that impacts of precipitation evenness persisted beyond systems that are generally defined as water-limited (Fig. 3). Our approach used site-specific means and variability to quantify highly even versus highly uneven precipitation. Thus, one standard deviation in precipitation evenness translated into similar impacts on plant biomass regardless of site, its typical precipitation evenness, or MAP (Fig. 2). While we show that plant growth declined in sensitivity to inter-annual variation in precipitation amount at sites with higher MAP (Fig. 3), sensitivity of plant growth to precipitation evenness did not diminish with higher MAP. These findings support a growing body of evidence that plant systems adapted to abundant annual precipitation amounts are still vulnerable to shifts in precipitation evenness due to impactful temporal periods of dry conditions and high evaporative demands ^19,24^. Wetter locations have plant communities with different functional traits than dry areas, such as larger leaves and taller plant height that increase primary productivity but may also create a sensitivity to periods with low water availability. For example, plants in wet and dry ecosystems have different critical moisture thresholds ^15,30^. This may explain, in part, why precipitation evenness can have sizeable impacts on plant biomass in systems with high MAP that are also less tolerant of soil drydown ^30^ and generally have high leaf area that enhance sensitivity to high evapotranspiration.

Benefits of less even precipitation for plant growth during dry years may be magnified in locations where warmer temperatures reduce the efficacy of small rainfall events due to high evaporation^25^. We tested whether the interactive effect of precipitation evenness and annual amount persisted across systems with warm and cold wet seasons. We found that high precipitation evenness continued to amplify the impacts of wet and dry years on plant growth regardless of the temperature of the wettest quarter (t = 0.18, p = 0.86)(Fig. S3), but that precipitation evenness has an additional direct positive effect on plant growth in warm systems regardless of annual precipitation amount (t = 2.18, p = 0.03). Precipitation evenness may have had a larger impact on plant growth in locations with warmer wet seasons due to the increased likelihood that plants are actively growing during precipitation and able to respond to large pulses or the lack thereof. Moreover, higher temperatures will exacerbate water limitations for plants by accelerating soil drying between rainfall events via increased vapor pressure deficit ^31^.

While discussion of decreases in precipitation evenness is relatively common across climate change studies, much less attention has been given to understanding the impacts of more even precipitation. We show that more frequent small storms (high evenness) have large impacts on plant biomass by amplifying the impacts of dry and wet years on plant growth and thereby increasing the variability in plant biomass (Fig. 2). Our results indicate that a broader perspective and awareness regarding both less and more even precipitation is needed as historic trends show that precipitation evenness is not universally declining, with many locations experiencing more even precipitation (Fig. 4). Projections indicate that emission scenarios may determine the intensity and scale of future changes in precipitation evenness ^1,32^.

**Figure 4.**
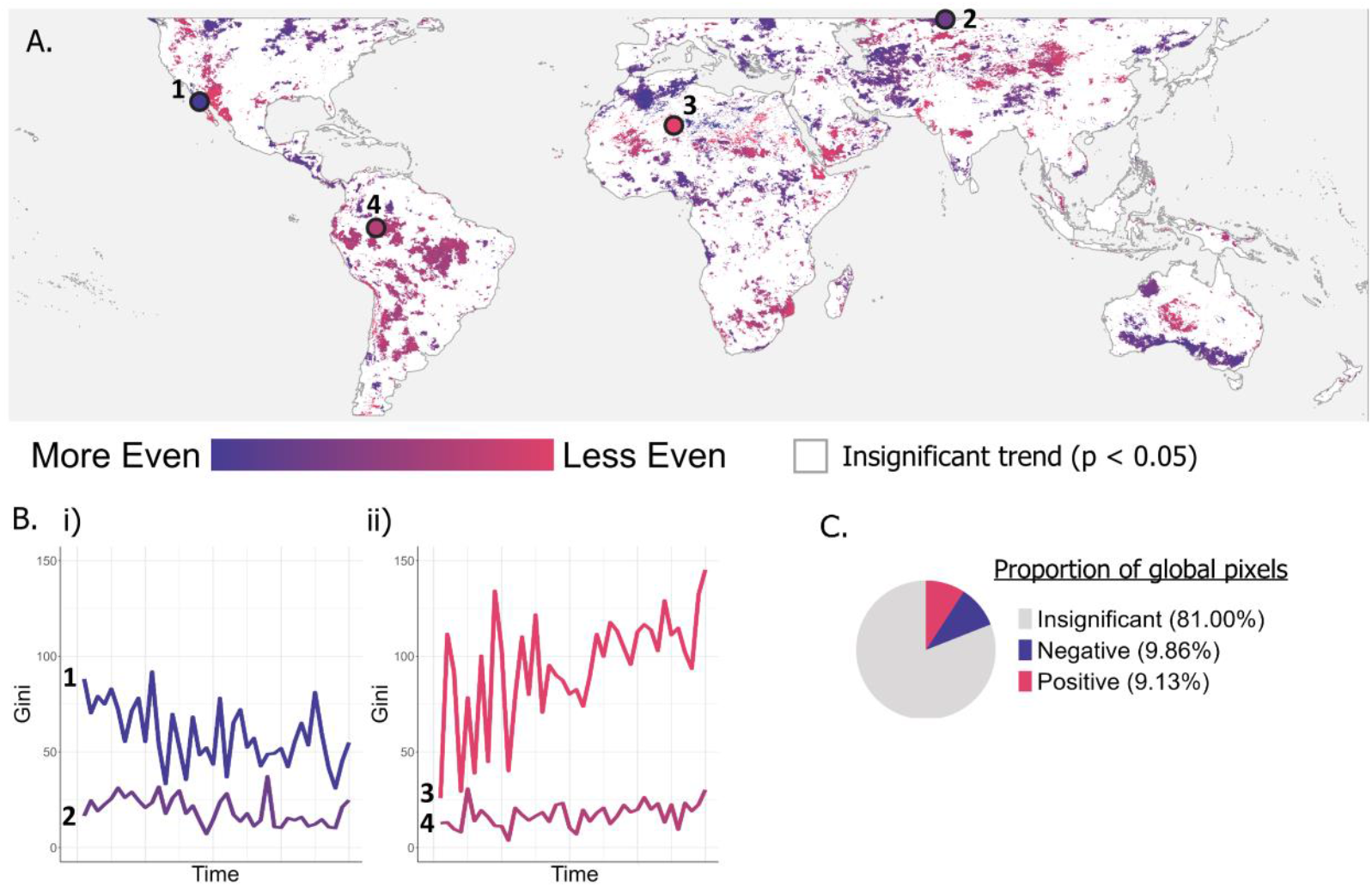
Historic temporal trends in precipitation evenness (1984 – 2023). Global (50°S – 50°N) spatial patterns in the temporal trend of precipitation evenness. Precipitation evenness is calculated as the unranked Gini coefficient for intra-annual precipitation patterns. Slope is indicated by Sen ‘s slope of the time series (Panel A). Areas with a statistically significant trend (Mann-Kendall, p < 0.05) are colored according to the gradient, land areas with an insignificant trend are shown in white. Four points show time series graphs of locations with i) significantly decreasing and ii) significantly increasing trends in UGI over 40 years. C. the proportion of pixels with insignificant, negative and positive trends in UGI.

Plant biomass was best explained by models that used precipitation metrics from the current water year until peak biomass. Models that used precipitation metrics calculated from shorter periods (60- and 30-day periods preceding peak biomass) were less able to explain interannual plant biomass (ΔAICc > 5, Table S4) and indicated that impacts of precipitation are determined more by amount independent of evenness. We anticipate that precipitation metrics from longer precipitation time frames were more informative for plant biomass as they capture accumulated impacts of precipitation on water and plants. While biotic legacy effects of previous years ‘ precipitation are frequently discussed ^33–35^, understanding the temporal consistency of preceding precipitation within the year (evenness) may also determine responsivity to future precipitation events within the same year. Using shorter time-frames for precipitation metrics likely removes these effects and focuses more on the immediate quantity of water for growth.

### Evenness Interaction with Nutrient Limitation

The addition of N, P, and K (all combined) did not alter the effect of precipitation evenness on plant biomass (t = -0.03, p = 0.98), nor did it affect the interaction between precipitation evenness and annual precipitation amount (t = 0.04, p = 0.97). This indicates that, overall, the effects of precipitation evenness on plant biomass are not driven by impacts on nutrient limitation (Fig. 2). Previous studies have hinted that precipitation evenness may regulate nutrient availability through the effect of inconsistent soil moisture that accompanies uneven precipitation ^16–18,22^, but the links to plant growth are inconsistent ^22,36^. More drying-rewetting cycles can increase nitrifier populations ^37^, and create periods when plant absorption is reduced more than N and P mineralization ^7,38,39^. Following an increase in soil nutrients, rewetting can create short periods of favorable growing conditions when neither water nor nutrients are limiting ^7^, or potentially increase leaching of nutrients ^40^. Our results indicate that the impacts of precipitation evenness are driven by water balance more than effects on nutrient limitations (Fig. 2). While we anticipate that less even precipitation may increase nutrient availability, it is also possible that the benefit of any additional soil nutrients is negated with growth being further limited to shorter periods of water availability ^7,41^.

### Precipitation Evenness Trends

Using a Sens ‘ slope estimate and Mann-Kendall tests across a 40-year time series for every pixel in the CHIRPS dataset, we found that UGI (precipitation unevenness) significantly increased and decreased across 9.1% and 9.8% of pixels respectively (1984-2023, Fig. 4). Precipitation is projected to become more uneven due to a warming atmosphere with the projected changes under current emissions scenarios exceeding those observed during the past 40 years ^1^. However, our results suggest that increasing evenness of precipitation is as widespread a phenomenon as increasing unevenness in terrestrial regions worldwide (Fig 4. A, C). This is a particularly important result given that high precipitation evenness amplifies the effect of anomalous annual rainfall on plant biomass, both positive and negative (Fig. 2, 3). Areas of increasing evenness and unevenness are distributed worldwide, demonstrating the global implications of our results. Globally, both pixel-based and site-based analyses showed that precipitation evenness is highly variable year-on-year (Fig. 2, 4B) and was not directly correlated to inter-annual annual precipitation variability (N = 537, R_spearman_ = 0.01) or MAP (N = 537, R_spearman_ = 0.02). This result indicates that the impacts of unusually even or uneven precipitation years could be felt across the globe, including in areas without a significant overall trend.

## Methods

We use annual biomass measurements within control and nutrient addition plots at 48 field sites globally (Table S1, Fig. S2) along with daily precipitation data to quantify the independent and interactive effects of annual precipitation amounts and intra-annual precipitation evenness on plant growth. We used nutrient addition plots within our models to test if the impact of precipitation evenness diminishes in plots with minimized nutrient limitation. We also mapped temporal trends in precipitation evenness across the globe, to identify regions experiencing significant increases or decreases in precipitation evenness.

### Precipitation Data

Our approach used daily rainfall data from the CHIRPS daily rainfall dataset v2.0 ^42^. This dataset was chosen for its large spatial coverage (all terrestrial areas between 50°S and 50°N) and fine spatial resolution (1 km). CHIRPS daily rainfall data is a gridded precipitation dataset that comes from both weather station data and satellite radar precipitation estimates. It has been used in similar studies and is shown to approximate amounts well but may have decreased accuracy in locations with few weather stations ^43^. To improve confidence in the use of CHIRPS data, we compared our global trend analysis (Fig. 4) with another study ^32^ which used a similar Gini metric approach (without linking to plant growth) based on data from 12,513 weather stations; both approaches found similar global patterns in precipitation evenness trends. To focus on relevant temporal precipitation inputs for plant ecosystems, we retrieved rainfall data for the water year beginning on October 1^st^ in the northern hemisphere, and July 1^st^ in the southern hemisphere. For the site-level analysis, the water year ended at the harvest date (peak biomass) for each site. For the analysis of global trends in precipitation without regard to plant growth, the water year was one calendar year. We also analyzed the sensitivity of our results to the selected time frame by comparing model output between our water year model (Model 1) and models that use precipitation data solely from the 30 and 60 days preceding peak biomass date (harvest). All precipitation data, both for sites and for global trends analysis, were obtained using Google Earth Engine ^44^.

### Evenness Metric

Our precipitation evenness metric is an unranked Gini index for a time series of precipitation data covering the water year, a standard timeframe for hydrological modelling that captures the natural progression of hydrologic seasons. This temporal period was chosen as it captures hydrological inputs prior to the growing season that strongly influence water availability within the growing season. Moreover, using a longer time frame (relative to the growing season) ensures that the research question remains focused on intra-annual precipitation evenness rather than singular events of drought and deluge.

To calculate the Gini index, daily precipitation is sorted by date (start of water year to harvest date, or to end of the water year) and summed cumulatively across time as a percentage of total precipitation for the total time period, thus forming a “Lorenz” curve (Fig. 1). The Gini index is calculated by summing the absolute area between the 45° line (representing a uniform/perfectly even precipitation distribution) and the Lorenz curve for each year (Fig. 1). High Gini values indicate a departure from an even temporal distribution and reflect a shift in temporal precipitation variability that may not be clear using other precipitation metrics (Fig. 1). This approach has been used in previous analyses, but either as a spatial variable ^24^, a ranked Gini index ^32^, or at small scales ^45,46^.

### Vegetation data

Plant biomass measurements estimates come from 48 grassland field sites of diverse climates and soils (Table S1). Each site contains three experimental blocks with both control plots and plots with a combined nutrient addition treatment of Nitrogen, Phosphorus, and Potassium ^47^. Plot size was 5 x 5 meters. Biomass at each site was clipped from a randomly selected, but permanent subplot (1×1 m) within each plot. Biomass clipping occurred at ground level within two 1 m × 0.1 m strips (totaling 0.2 m^2^) and was scaled to a square meter basis.

Sampling occurred in a spatial sequence that avoided resampling of plants previously clipped. Dead biomass was discarded, and live plant biomass was dried in an oven at 60°C for 24 hours, and weighed to the nearest 0.01 g. If sampling occurred more than once per year, as is the case at two of our sites with bimodal growth patterns, the annual sum of biomass for the two sample dates was used. In our analysis, plant live biomass from either control plots or nutrient addition plots were averaged across blocks for each year, to produce one value for each site in each year.

### Statistical Approach

We employed a linear model in R ^48^ to test the effect of annual precipitation amount on site-level live biomass under scenarios of high, average, and low precipitation evenness visualize in Figure 2. Annual precipitation amount and precipitation evenness were scaled using a site-specific mean and standard deviation (2004-2023) to create annual anomaly variables. Site level biomass was calculated by scaling each annual value against the site-specific mean and standard deviation. Observations were binned according to annual scaled precipitation evenness; high and low evenness years were considered as annual scaled values one standard deviation greater or less than the 20-year mean (2003-2024). This model used site-relative biomass as the response variable and scaled annual precipitation amount as the explanatory variable. In order to visualize the individual and combined impact of precipitation amount and evenness on plant biomass (H1 and H2), the models were repeated using plant biomass measures from both control and nutrient addition plots.

Next, we used a linear mixed model ^49^ in R ^48^ (Model 1) to quantify the impact of precipitation evenness (UGI) on annual live plant biomass (*Biomass*) using control (no nutrient addition) plots (H1). We tested for a direct effect of interannual meterics of precipitation evenness (*B*_*3*_) as well as an interactive effect (*B*_*5*_) of precipitation evenness and annual precipitation amount anomalies (*Pdev*). To test if impacts of precipitation evenness differed at sites with low and high average annual precipitation amounts, we used a three-way interaction (*B*_*7*_) of precipitation evenness, annual precipitation anomalies (scaled), and mean annual precipitation amount (*MAP*). We included temperature of wettest quarter (*TWQ*) to account for variation in evaporative demands and a random effect of site identity (*Site*) to account for important non-precipitation spatial factors that have large impacts on plant resource availability. A variance inflation factor (VIF) was calculated using the car package ^50^ to ensure that none of our explanatory variables in our models were co-linear, with all explanatory variables producing VIF values less than 1.1. We used a normal error distribution. The results of Model 1 for biomass are shown in Fig. 3.

Model 1

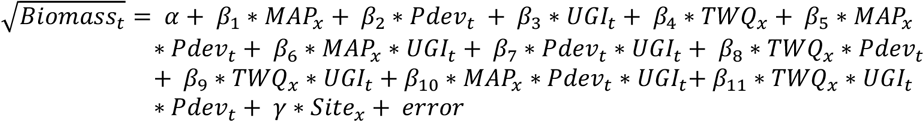

We added coefficients to a subset of Model 1 to create Model 2, a model that allows us to test if the effect of precipitation evenness differs in control versus nutrient addition plots (*NPK*) where nutrients are not limiting (H2). Model 2 uses data from both control and nutrient addition plots (N = 1111). We tested for both an interactive effect of nutrient addition and precipitation evenness (*B*_*6*_) and an interactive effect of nutrient addition with annual precipitation anomalies and precipitation evenness (*B*_*7*_).

Model 2

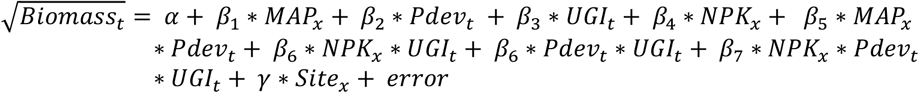

All models were tested to ensure that residuals were normally distributed and heteroskedastic using diagnostic plots from the performance package ^51^ (Fig. S4-S8). All models were checked against similar models, but with the evenness terms removed, and found to improve model fit using the log-likelihood metric. We tested our models for spatial autocorrelation using a semivariogram on the residuals of our model before and after adding a site identity random effect. This analysis revealed that the apparent spatial autocorrelation in a fixed-effects-only model was eliminated by the addition of a site identity random effect (Fig S8).

We evaluated temporal autocorrelation by calculating the mean lag-1 autocorrelation of model residuals across sites. We found a low value (φ = 0.0843), indicating weak serial dependence. Residual ACF plots confirmed that correlations at successive time steps were minimal, suggesting that temporal autocorrelation was not a concern in this dataset (Fig S9).

To investigate global trends in precipitation evenness over time, we calculated the Gini index for every CHIRPS pixel (terrestrial pixels 50°S - 50°N, using the appropriate water year window for the southern and northern hemispheres) for every year 1984 – 2023, to create 40-year time series. Sen ‘s slope was calculated to estimate the magnitude of the trend over time, and Mann-Kendall tests to assess the statistical significance of the trend per 1×1km pixel.

Global distribution and proportion of significant increases and decreases in precipitation evenness are shown in Figure 4.

## Acknowledgements

We would like to thank the many data contributors from the Nutrient Network who contributed data but were not listed as authors (Table S3). We would also like to thank the editors and anonymous reviewers for their efforts to improve our manuscript. Views and opinions expressed are however those of the author(s) only and do not necessarily reflect those of the European Union or the European Research Council. Neither the European Union nor the granting authority can be held responsible for them.

## Author Contributions

Conceptualization: TT, SH, AJ

Visualization: TT, AH, SH

Funding acquisition: EB, ES, AJ

Investigation: All authors (except SH, QC, and AH)

Writing – original draft: TT, SH, AH, QC

Writing – review & editing: All authors

Competing interests: The authors have no competing interests to declare

## Supplementary Materials

Figs. S1 to S9

Tables S1 to S4

