## Supplementary Materials for "Intra-annual precipitation variability mutes or magnifies the impact of wet and dry years on plant biomass"

### Intra-annual precipitation patterns mute or magnify the impact of wet and dry years on plant biomass.

**File includes:**
**Figs. S1 to S9**
**Tables S1 to S3**

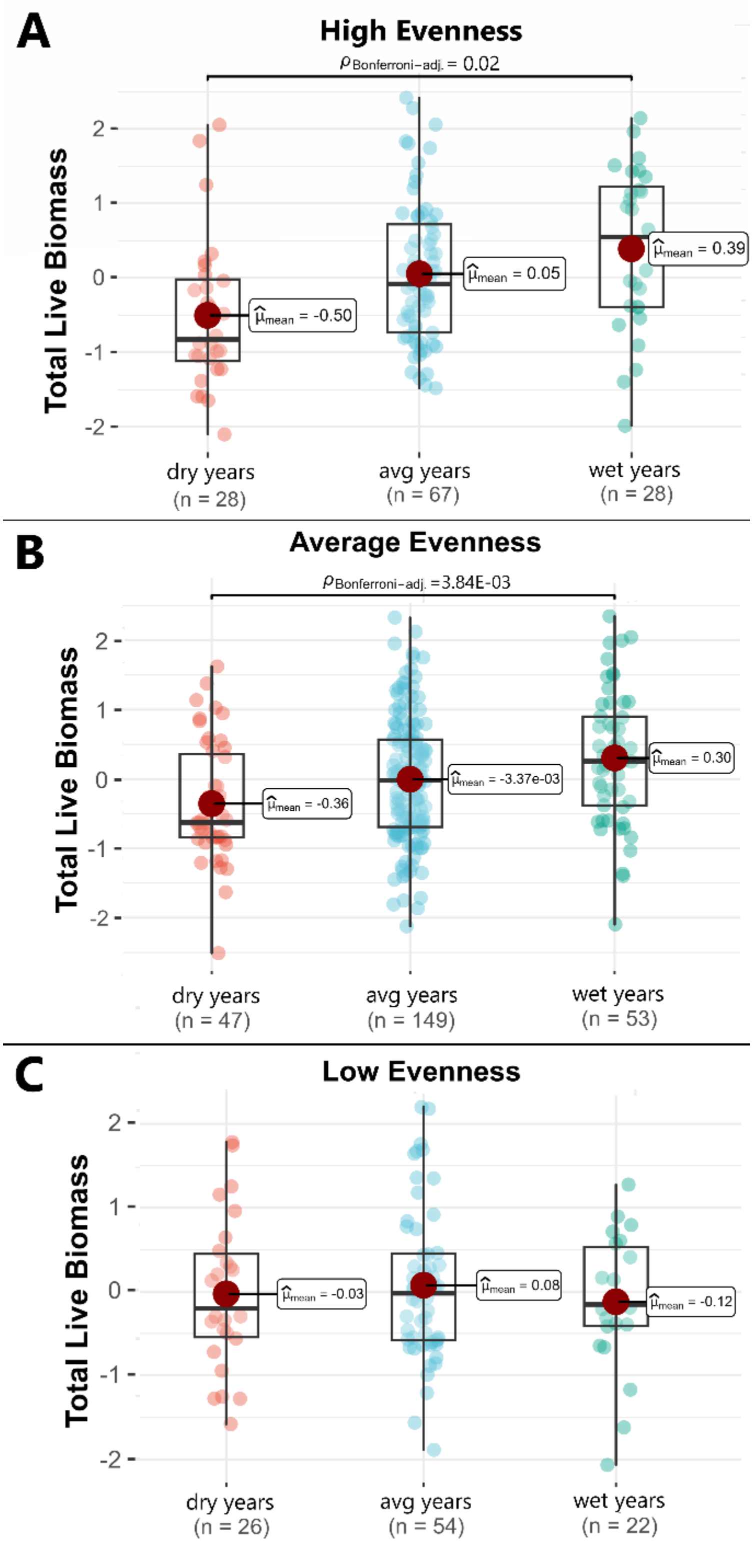


**Fig. S1. Average impacts of wet and dry years on plant biomass are only apparent under highly even intra-annual precipitation patterns.** Points represent total live biomass relative to site means under high, average, and low precipitation evenness scenarios. Dry and wet year scenarios are defined as annual amounts of one standard deviation above or below the site-specific average (2004-2023). High and low evenness scenarios represent values greater than one standard deviation above or below the site-specific evenness average (2004-2023). Statistics bar above boxplots indicates significant differences using a pairwise comparison and the respective Bonferroni-corrected p value.

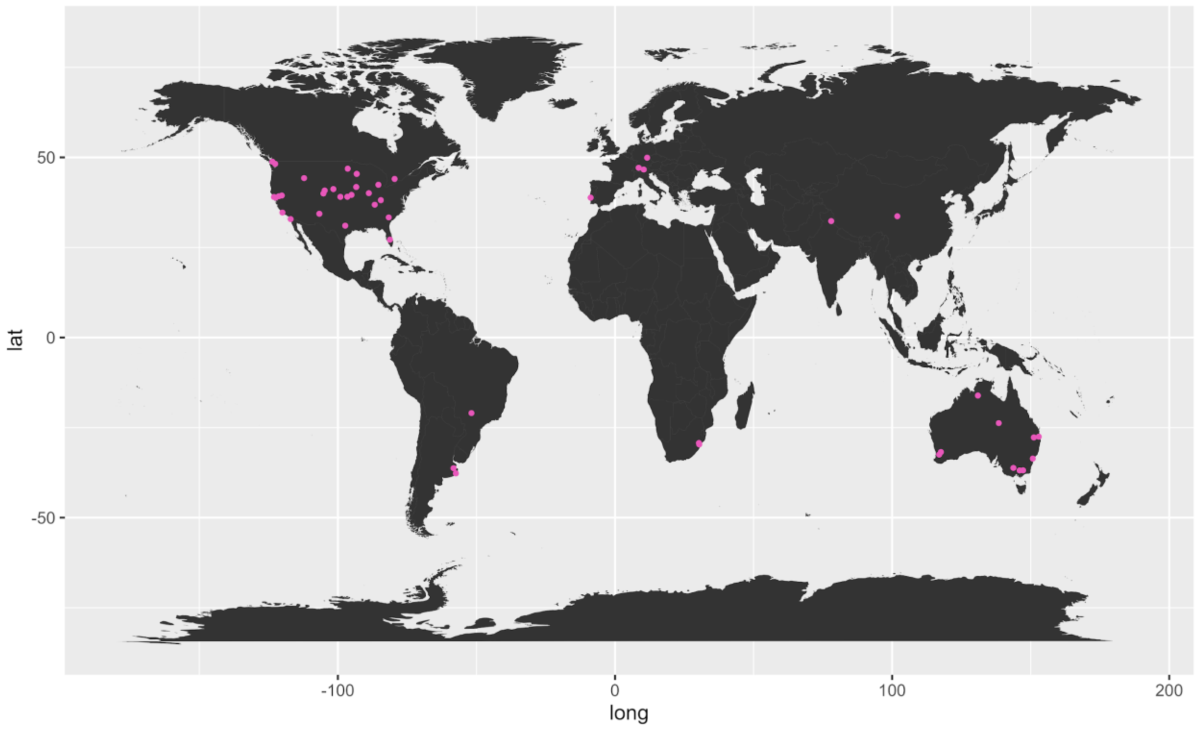


**Fig. S2. Map of study site locations.** Pink dots represent individual experimental sites from the Nutrient Network that were included in this study.

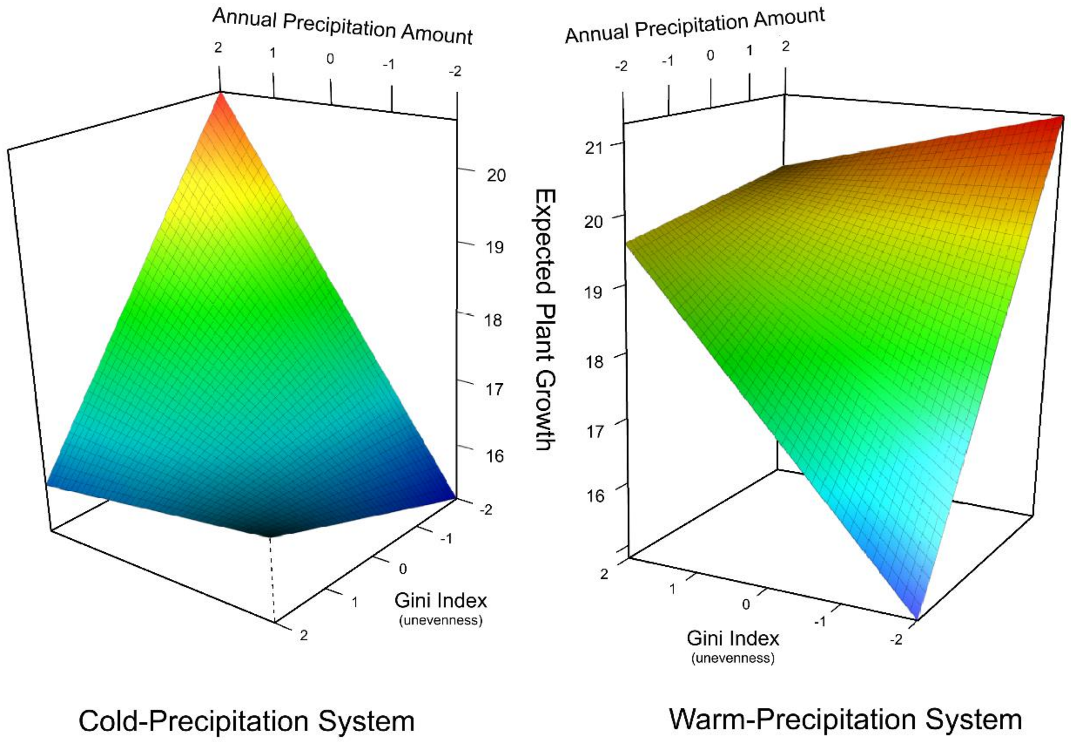


**Fig. S3. Predicted plant growth based on model predictions for systems that receive the majority of their precipitation in colder (left) and warmer (right) temperatures.** Gini index indicates the evenness of the precipitation relative to the 20-year mean and standard deviation. Annual amount is the amount of annual precipitation relative to the 20-year mean and standard deviation. Expected plant growth is the predicted amount of live plant biomass based on model predictions (R square = 0.73).

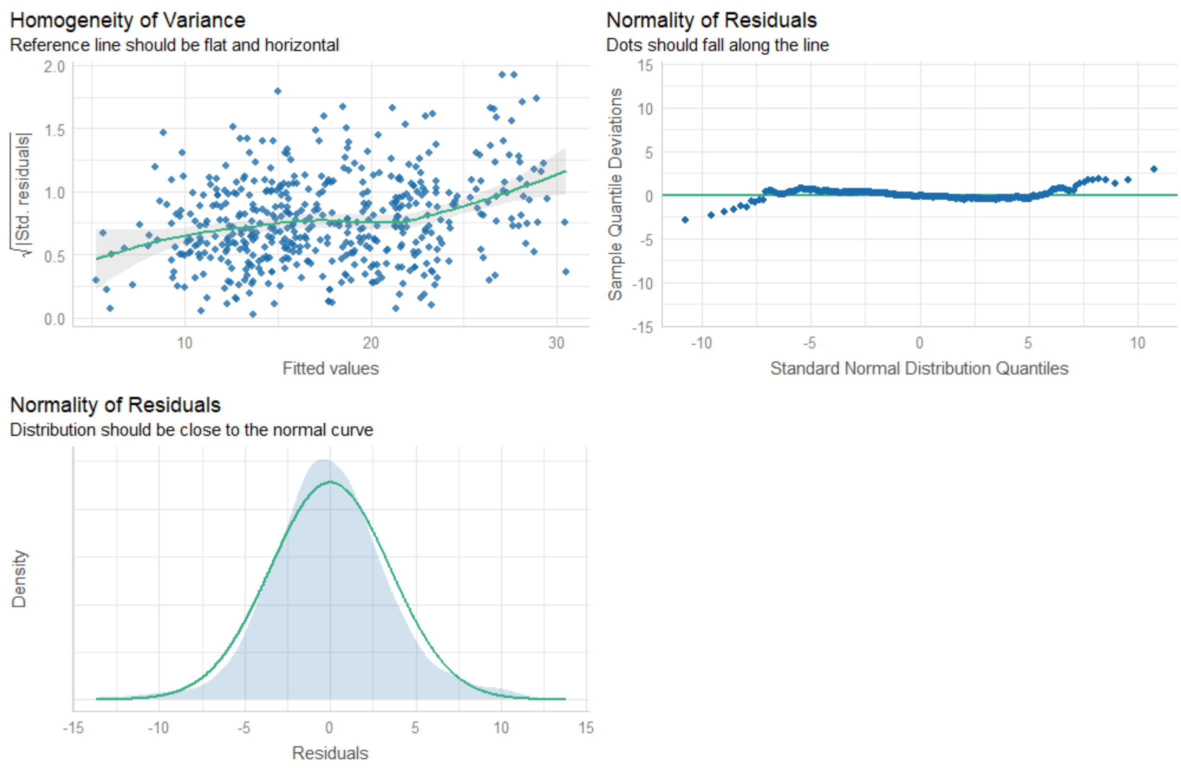


**Fig. S4. Diagnostic Plots for Model 1.**

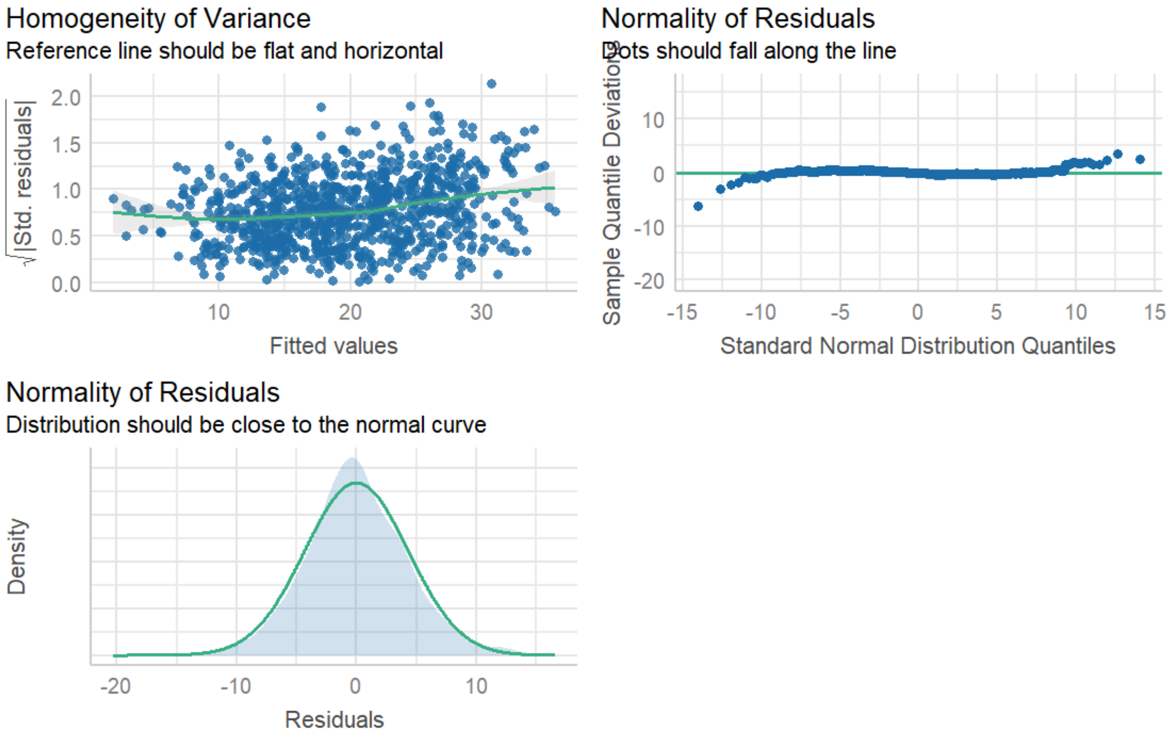


**Fig. S5. Diagnostic Plots for Model 2.**

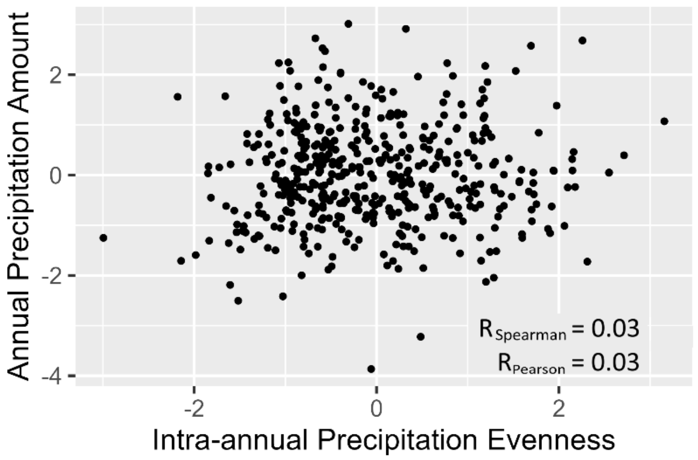


**Fig. S6. Correlation plot for precipitation evenness vs annual precipitation amounts.** The annual precipitation amounts and precipitation evenness are scaled values indicating deviation from site-specific means.

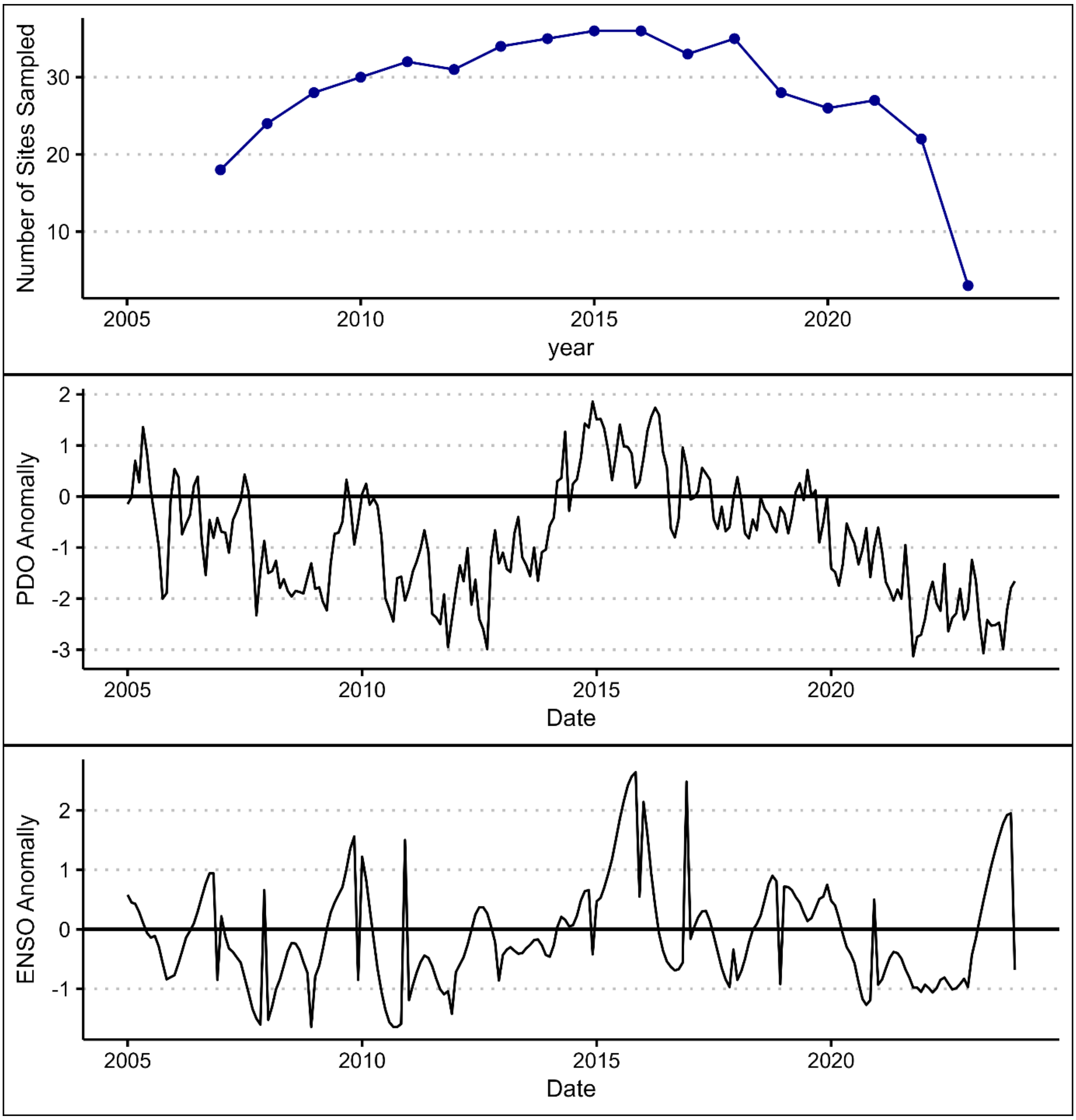


**Fig. S7. Temporal sampling distribution and relevant climate cycles.** The top panel indicates the number of sites sampled each year. The middle and lower panels indicate the co-occurring climate cycles of the Pacific Decadal Oscillation (PDO) and El Nino/Southern Oscillation (ENSO) respectively. PDO data is graphed using monthly data, and ENSO data represents anomalies of the ENSO index (using three month averages in Nino 3.4). All climate data comes from NOAA's extended reconstruction of sea surface temperatures (ERSST v5).

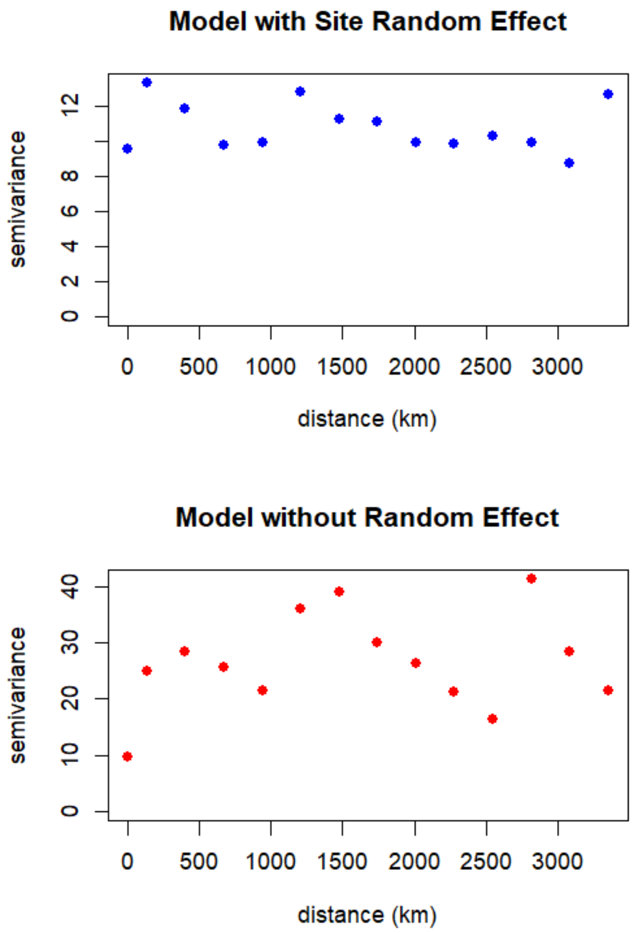


**Fig. S8. Semivariograms comparing spatial autocorrelation of models with and without a site-identity random effect.** The top panel displays the semivariance of residuals at sites of various distances when using a model that includes site name as a random effect. The bottom panel displays the same relationship but uses a model without a random effect.

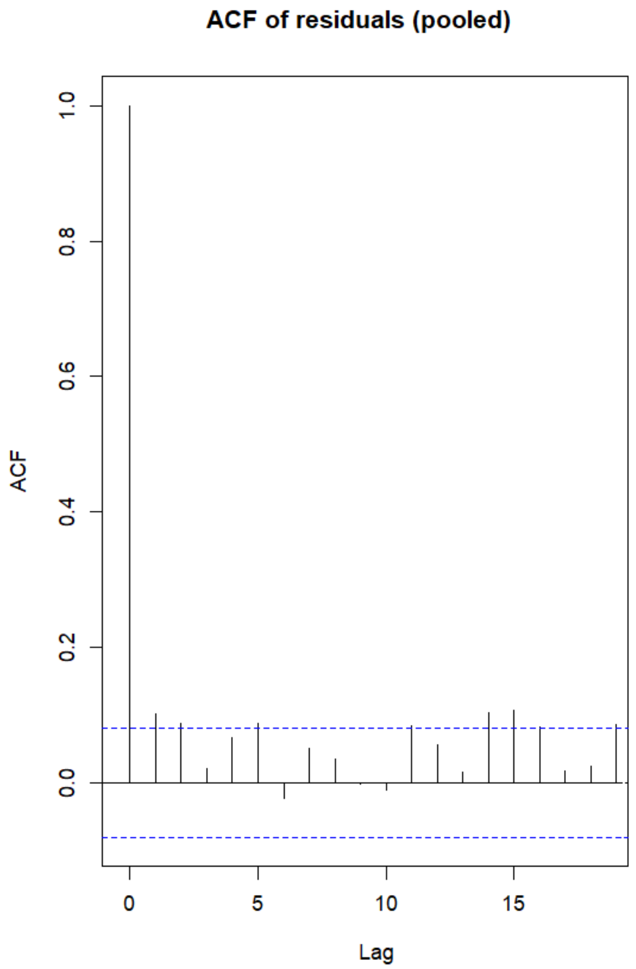


**Fig. S9. Autocorrelation values (ACF) of residuals at different time lags.**

**Table S1. Table of site locations and climatic characteristics.** Sites come from the Nutrient Network and include various habitat types. Abbreviations include aridity index (AI), potential evapotranspiration (PET), mean annual temperature (MAT), and mean annual precipitation (MAP).

| Site Name | Habitat | Elevation (m) | Latitude | Longitude | AI | PET | MAT (°C) | MAP (mm) |
|---|---|---|---|---|---|---|---|---|
| Archbold Biological Station | mixed grass prairie | 8 | 27.17 | -81.22 | 0.68 | 1784 | 22.7 | 1314 |
| Azi | alpine grassland | 3500 | 33.67 | 101.87 | 0.72 | 990 | 1.4 | 745 |
| Bayreuth | mesic grassland | 340 | 49.92 | 11.58 | 0.91 | 828 | 8.5 | 522 |
| Benedictine Bottoms | tallgrass prairie | 240 | 39.60 | -95.09 | 0.66 | 1435 | 12.4 | 912 |
| Bogong | alpine grassland | 1760 | -36.87 | 147.25 | 1.59 | 1056 | 6.0 | 1392 |
| Boulder South Campus | shortgrass prairie | 1633 | 39.97 | -105.23 | 0.28 | 1731 | 9.9 | 476 |
| Burrawan | semiarid grassland | 425 | -27.74 | 151.14 | 0.34 | 1872 | 18.2 | 584 |
| Cedar Creek LTER | tallgrass prairie | 270 | 45.43 | -93.21 | 0.71 | 1044 | 6.3 | 777 |
| Cedar Point Biological Station | shortgrass prairie | 965 | 41.21 | -101.64 | 0.28 | 1615 | 9.6 | 492 |
| Chichaqua Bottoms | tallgrass prairie | 274 | 41.79 | -93.39 | 0.72 | 1202 | 9.3 | 940 |
| Companhia das Lezirias | annual grassland | 20 | 38.83 | -8.79 | 0.47 | 1481 | 16.6 | 637 |
| Cowichan | old field | 50 | 48.81 | -123.63 | 1.21 | 918 | 10.4 | 1072 |
| Elliott Chaparral | annual grassland | 200 | 32.88 | -117.05 | 0.18 | 1868 | 17.7 | 211 |
| Ethabuka (Main Camp) | desert grassland | 104 | -23.76 | 138.47 | 0.06 | 2980 | 24.1 | 166 |
| Fruebuel | pasture | 995 | 47.11 | 8.54 | 1.76 | 869 | 7.0 | 1354 |
| Hall's Prairie | tallgrass prairie | 194 | 36.87 | -86.70 | 0.97 | 1335 | 13.8 | 1334 |
| Hopland REC | annual grassland | 598 | 39.01 | -123.06 | 0.66 | 1608 | 13.2 | 846 |
| Kellogg Biological Station LTER | old field | 288 | 42.41 | -85.39 | 0.82 | 1102 | 8.8 | 957 |
| Kibber (Spiti) | alpine grassland | 4241 | 32.32 | 78.01 | 0.33 | 1198 | -1.5 | 264 |
| Kidman Springs | savanna | 87 | -16.11 | 130.95 | 0.28 | 2611 | 27.3 | 745 |
| Kinypanial | semiarid grassland | 90 | -36.20 | 143.75 | 0.22 | 1851 | 15.6 | 395 |
| Koffler Scientific Reserve | pasture | 301 | 44.02 | -79.54 | 0.96 | 892 | 6.3 | 923 |
| Konza LTER | tallgrass prairie | 440 | 39.07 | -96.58 | 0.58 | 1542 | 12.1 | 891 |
| Las Chilcas | mesic grassland | 15 | -36.28 | -58.27 | 0.72 | 1334 | 15.1 | 964 |
| Mar Chiquita | grassland | 6 | -37.72 | -57.42 | 0.70 | 1302 | 14.3 | 926 |

| | | | | | | | | |
|---|---|---|---|---|---|---|---|---|
| Mclaughlin UCNRS | annual grassland | 642 | 38.86 | -122.41 | 0.54 | 1719 | 14.0 | 629 |
| Minnesota State University | tallgrass prairie | 311 | 46.87 | -96.45 | 0.52 | 1071 | 5.0 | 623 |
| Mt Gilboa | montane grassland | 1748 | -29.28 | 30.29 | 0.63 | 1492 | 14.1 | 810 |
| Mt. Caroline | savanna | 285 | -31.78 | 117.61 | 0.15 | 2188 | 17.7 | 296 |
| Nillahcootie | old field | 280 | -36.90 | 146.01 | 0.63 | 1503 | 13.8 | 869 |
| Pingelly Paddock | old field | 338 | -32.50 | 116.97 | 0.22 | 2052 | 16.3 | 433 |
| Pinjarra Hills | pasture | 38 | -27.53 | 152.92 | 0.59 | 1825 | 20.0 | 1000 |
| Sagehen Creek UCNRS | montane grassland | 1920 | 39.43 | -120.24 | 0.57 | 1469 | 5.8 | 930 |
| Saline Experimental Range | mixed grass prairie | 555 | 39.05 | -99.10 | 0.35 | 1745 | 12.1 | 640 |
| Savannah River | savanna | 71 | 33.34 | -81.65 | 0.76 | 1556 | 17.4 | 1211 |
| Sedgwick Reserve UCNRS | annual grassland | 550 | 34.70 | -120.02 | 0.25 | 1890 | 15.6 | 347 |
| Sevilleta LTER | desert grassland | 1600 | 34.36 | -106.69 | 0.12 | 2166 | 13.1 | 231 |
| Sheep Experimental Station | shrub steppe | 1661 | 44.26 | -112.21 | 0.18 | 1336 | 5.3 | 230 |
| Shortgrass Steppe LTER | shortgrass prairie | 1650 | 40.82 | -104.77 | 0.21 | 1718 | 8.9 | 337 |
| Sierra Foothills REC | annual grassland | 197 | 39.24 | -121.28 | 0.49 | 1909 | 16.3 | 634 |
| Smith Prairie | mesic grassland | 63 | 48.21 | -122.62 | 0.65 | 926 | 10.2 | 557 |
| Spindletop | pasture | 271 | 38.13 | -84.50 | 0.91 | 1266 | 12.5 | 1233 |
| Temple | tallgrass prairie | 184 | 31.04 | -97.35 | 0.47 | 1862 | 19.4 | 861 |
| Trelease | tallgrass prairie | 200 | 40.08 | -88.83 | 0.80 | 1233 | 11.1 | 993 |
| Tres Lagoas | cerrado | 279 | -20.98 | -51.80 | 0.69 | 1666 | 23.2 | 1129 |
| Ukulinga | mesic grassland | 842 | -29.67 | 30.40 | 0.53 | 1560 | 17.7 | 721 |
| Val Mustair | alpine grassland | 2320 | 46.63 | 10.37 | 0.93 | 731 | 0.1 | 628 |
| Yarramundi | mesic grassland | 19 | -33.61 | 150.74 | 0.52 | 1630 | 17.3 | 744 |

**Table S2. Table of coauthors and their contributions.**

| Last Name | First Name | Developed and framed research question(s) | Analyzed data or generated new dataset | Contributed to data analyses | Wrote the paper | Contributed to paper writing | Current site- level coordinator | Current network-level coordinator |
|---|---|---|---|---|---|---|---|---|
| Terry | Tyson J. | x | x | x | x | | | |
| Higgins | Steven I. | x | | x | | x | | |
| Hammer | Alexandra | x | x | x | | | | |
| Jentsch | Anke | | | | | x | x | |
| Chen | Qingqing | x | | | | x | | |
| Risch | Anita C. | | | | | x | x | |
| Hagenah | Nicole | | | | | x | x | |
| Collins | Scott | | | | | x | x | |
| Raynaud | Xavier | | | | | x | x | |
| Yost | Jennifer | | | | | x | x | |
| Seabloom | Eric W. | | | | | x | x | x |
| Borer | Elizabeth T. | | | | | x | x | x |
| Adler | Peter B. | | | | | x | x | |
| Dickman | Chris R. | | | | | x | x | |
| Brudvig | Lars A. | | | | | x | x | |
| Caldeira | Maria C. | | | | | x | x | |
| Knops | Johannes | | | | | x | x | |
| Niu | Yujie | x | | | | x | | |
| Komatsu | Kim | | | | | x | x | |
| Wardle | Glenda | | | | | x | x | |
| Catford | Jane | | | | | x | x | |
| Bugalho | Miguel | | | | | x | x | |
| Bakker | Jonathan D. | | | | | x | x | |

**Table S3. Table of data contributors not listed as authors.**

| Site Name | Last Name | First Name | PI Start Year | PI End Year |
|---|---|---|---|---|
| Archbold Biological Station | Boughton | Elizabeth | 2015 | |
| Azi | Chu | Chengjin | 2007 | 2012 |
| Azi | Du | Guozhen | 2007 | 2012 |
| Azi | Li | Qi | 2007 | 2012 |
| Azi | Li | Wei | 2007 | 2012 |
| Azi | Wen | Gang | 2007 | 2012 |
| Boulder South Campus | Davies | Kendi | 2008 | 2016 |
| Boulder South Campus | Melbourne | Brett | 2008 | 2016 |
| Benedictine Bottoms | Mortensen | Brent | 2017 | |
| Benedictine Bottoms | Paper | Janet | 2017 | |
| Bogong | Moore | Joslin | 2009 | |
| Bogong | Morgan | John | 2009 | |
| Burrawan | Buckley | Yvonne | 2008 | 2012 |
| Burrawan | Firn | Jennifer | 2008 | 2019 |
| Chichaqua Bottoms | Biederman | Lori | 2009 | |
| Chichaqua Bottoms | Harpole | W. | 2009 | 2014 |
| Chichaqua Bottoms | Hofmockel | Kirsten | 2009 | 2014 |
| Chichaqua Bottoms | Sullivan | Lauren | 2009 | 2014 |
| Cedar Creek LTER | Harpole | W. | 2007 | 2014 |
| Cedar Creek LTER | Kay | Adam | 2007 | 2010 |
| Cedar Point Biological Station | Knops | Johannes | 2007 | |
| Cedar Point Biological Station | Wheeler | George | 2018 | |
| Las Chilcas | Chaneton | Enrique | 2013 | 2019 |
| Las Chilcas | Tognetti | Pedro | 2013 | |
| Las Chilcas | Yahdjian | Laura | 2013 | |
| Cowichan | MacDougall | Andrew | 2007 | |
| Elliott Chaparral | Cleland | Elsa | 2008 | |
| Fruebuel | Güsewell | Sabine | 2008 | 2016 |
| Fruebuel | Hautier | Yann | 2008 | 2016 |
| Fruebuel | Hector | Andy | 2008 | 2016 |
| Mt Gilboa | Kirkman | Kevin | 2010 | |
| Mt Gilboa | Tedder | Michelle | 2010 | |
| Hall's Prairie | McCulley | Rebecca | 2007 | 2014 |
| Hall's Prairie | Nelson | Jim | 2007 | 2014 |
| Hopland REC | Harpole | W. | 2007 | 2018 |
| Kibber (Spiti) | Sankaran | Mahesh | 2012 | |
| Kidman Springs | Richards | Anna | 2014 | 2023 |
| Kinypanial | Morgan | John | 2007 | |
| Koffler Scientific Reserve at Joker's Hill | Cadotte | Marc | 2010 | |
| Koffler Scientific Reserve at Joker's Hill | Weiss | Arthur | 2010 | |

| | | | | |
|---|---|---|---|---|
| Konza LTER | Smith | Melinda | 2007 | |
| Temple | Fay | Philip | 2007 | 2019 |
| Tres Lagoas | Lannes | Lucíola | 2015 | |
| Tres Lagoas | olde Venterink | Harry | 2015 | |
| Mar Chiquita | Alberti | Juan | 2012 | |
| Mar Chiquita | Daleo | Pedro | 2012 | |
| Mclaughlin UCNRS | Harpole | W. | 2007 | 2018 |
| Mt. Caroline | Prober | Suzanne | 2008 | |
| Pingelly Paddock | Price | Jodi | 2013 | 2021 |
| Pingelly Paddock | Standish | Rachel | 2013 | 2021 |
| Pinjarra Hills | Buckley | Yvonne | 2013 | 2014 |
| Pinjarra Hills | Dwyer | John | 2013 | |
| Sagehen Creek UCNRS | Gruner | Daniel | 2007 | 2013 |
| Sagehen Creek UCNRS | Yang | Louie | 2007 | 2013 |
| Saline Experimental Range | Smith | Melinda | 2007 | 2017 |
| Savannah River | Damschen | Ellen | 2007 | 2013 |
| Savannah River | Orrock | John | 2007 | 2013 |
| Sedgwick Reserve UCNRS | D'Antonio | Carla | 2007 | 2016 |
| Sedgwick Reserve UCNRS | Harpole | W. | 2007 | 2008 |
| Sevilleta LTER | Ladwig | Laura | 2007 | 2012 |
| Sevilleta LTER | Ohlert | Tim | 2019 | 2023 |
| Shortgrass Steppe LTER | Blumenthal | Dana | 2007 | 2022 |
| Shortgrass Steppe LTER | Brown | Cynthia | 2007 | 2022 |
| Shortgrass Steppe LTER | Klein | Julia | 2007 | 2022 |
| Shortgrass Steppe LTER | Knapp | Alan | 2007 | 2022 |
| Sierra Foothills REC | Harpole | W. | 2007 | 2018 |
| Smith Prairie | Hille Ris Lambers | Janneke | 2007 | 2013 |
| Spindletop | McCulley | Rebecca | 2007 | |
| Spindletop | Nelson | Jim | 2007 | |
| Temple | Martina | Jason | 2019 | |
| Temple | Yost | Jenifer | 2024 | |
| Trelease | Leakey | Andrew | 2008 | 2016 |
| Ukulinga | Hagenah | Nicole | 2009 | 2017 |
| Ukulinga | Kirkman | Kevin | 2009 | |
| Ukulinga | Tedder | Michelle | 2017 | |
| Val Mustair | Pichon | Noémie | 2024 | |
| Val Mustair | Risch | Anita | 2008 | |
| Val Mustair | Schuetz | Martin | 2008 | 2024 |
| Yarramundi | Ochoa Hueso | Raul | 2014 | |
| Yarramundi | Power | Sally | 2014 | |

**Table S4. Comparison of models and goodness of fit metrics.**

| **Model (precipitation window used)** | **Loglikelihood** | **AICc** | ΔAIC | **weight** |
|---|---|---|---|---|
| Water year-peak biomass | -3834.259 | 7691.0 | 0.00 | 0.935 |
| 60 days preceding peak biomass | -3836.980 | 7696.4 | 5.44 | 0.062 |
| 30 days preceding peak biomass | -3839.797 | 7702.0 | 11.08 | 0.004 |
